# Learning heritable multimodal brain representation via contrastive learning

**DOI:** 10.64898/2026.02.19.706893

**Authors:** Tian Xia, Xingzhong Zhao, Saiful Sheikh Muhammad Islam, Kamil Khan Mohammed, Ziqian Xie, Degui Zhi

## Abstract

Magnetic resonance imaging (MRI)-derived phenotypes (IDP) has enabled the discovery of numerous genomic loci associated with brain structure and function. However, most existing IDPs and learned representations are derived from a single imaging modality, missing complementary information across modalities and potentially limiting the scope of genetic discovery. Here, we introduce a multimodal contrastive learning framework to derive heritable representations from paired T1- and T2-weighted MRIs. Unlike single-modality reconstruction-based models, we designed a momentum-based contrastive learning framework. As a result, our approach offers improved prediction of traditional IDPs, age, and brain disorders. Notably, genome-wide association studies (GWAS) of the learned representations reveal a substantially higher overlap of genetic loci across modalities, indicating improved alignment of their underlying genetic architecture. Analysis of the GWAS loci identified shared protein and drug targets, yielding meaningful biological insights. Overall, our framework learns shared representations across brain imaging modalities that exhibit anatomical and genetic coherence.

## Introduction

Magnetic resonance imaging (MRI) is widely used to noninvasively characterize the structure and function of human organs, including the brain, heart, liver, and kidneys. Large-scale cohorts combining brain MRI and genotyping have established imaging-derived phenotypes (IDPs) as powerful quantitative traits for mapping the genetic architecture of the human brain structure, function and disease risk.

Researchers employed automated processing pipelines, such as FSL and FreeSurfer, to extract thousands of measures simultaneously across multiple MRI modalities, including structural, diffusion, and functional imaging. The first UK Biobank protocol reported approximately 4,350 IDPs [1], encompassing subcortical structure volumes, microstructural metrics in major tracts (e.g., DTI and NODDI measures), and structural and functional connectivity metrics. Leveraging these IDPs, a series of large-scale GWAS were conducted [2–6], identifying hundreds of genomic loci associated with diverse MRI-derived traits and revealing the genetic architecture underlying brain structure and function.

Recently, machine learning and deep learning approaches have emerged as unbiased methods for deriving heritable representations of brain structure directly from unlabeled MRI data, addressing limitations of conventional IDPs. Many commonly used IDPs, such as subregion volumes, are highly correlated and can be biased, as they rely on predefined brain segmentations for downstream analysis. Even widely used segmentation software exhibits biases and inconsistencies [7]. Consequently, methods that take raw brain images as input offer a promising alternative for defining MRI-based phenotypes.

A straightforward approach involves applying dimensionality reduction techniques, such as principal component analysis (PCA) or independent component analysis (ICA), directly to whole-brain images. ICA and its variants have long been used for feature extraction in functional MRI [8–10] and can also be extended to structural MRI [11, 12]. Other methods, such as non-negative matrix factorization (NMF), have been applied to reduce whole-brain imaging data into 32, 64, 128, or more patterns of structural covariation, termed patterns of structural covariation (PSC) [13]. Conceptually, these approaches can be viewed as linear analogs of autoencoders, as they learn low-dimensional representations while retaining the ability to reconstruct the original images.

Regarding deep learning approaches, our recent work employed reconstruction-based autoencoders to learn 128-dimensional representations of brain MRI, enabling the discovery of novel genetic loci associated with brain structure [14]. This study represented the first unsupervised learning-derived image endophenotype (UDIP). For clarity, we will name it Unsupervised Deep learning-derived Imaging Phenotype (UDIP), and we will refer all the IDP directly generated from FSL/FreeSurfer and reported by the UKBiobank as traditional IDP. Nevertheless, they all belong to the large concept of IDP. In the realm of semi-supervised learning, Wen et al. proposed a method to generate dimensional neuroimaging endophenotypes (DNE) using a framework called Smile-GAN (SeMI-supervised cLustEring-Generative Adversarial Network) [15]. This approach leverages disease labels to guide feature extraction, with a particular focus on capturing the heterogeneity of brain disorders. Several fully supervised learning methods have also been developed for MRI analysis [16–20]. Additionally, there are other computer vision studies applying deep learning to MRI data [21, 22], though these do not target imaging-genomic analyses and are therefore not detailed here.

Given all the methods mentioned above, the vast majority of existing IDPs or learned brain representations are constructed from a single imaging modality, inherently limiting the scope of information they can capture. This single-modal paradigm overlooks the fact that different MRI modalities probe complementary aspects of brain structure and tissue properties-for example, T1-weighted images emphasize macrostructural anatomy, whereas T2-weighted contrasts are sensitive to microstructural and pathological variations. As a result, representations learned from a single modality may be biased toward modality-specific features and fail to fully enable genetic discovery.

In this work, we introduce a **multimodal contrastive representation learning framework** [23] to learn common representations of brain structure across paired T1- and T2-weighted MRIs from the UK Biobank, followed by an analysis of the biological information encoded in these representations. Our work is based on the MoCoV2 (Momentum Contrast learning V2) [23] that is known to provide stable large sample training with limited batch size. In contrast to the original MoCoV2 framework, which forms positive pairs from different augmentations of the same image, we define positive pairs using multimodal MRI images acquired from the same individual. The architecture pulls representations of the same individual across modalities closer together while pushing representations of different individuals apart. Shared biological structure is then evaluated by comparing the overlap of genomic loci associated with representations learned from each modality. We name the learned common representation MM-UDIP (MultiModal MoCoV2 Unsupervised Derived Imaging Phenotypes). For comparison, we named the representation learned from our previous single-modal ViT-autoencoder-based method SM-UDIP (SingleModal ViT Unsupervised Derived Imaging Phenotypes) [14, 24].

Through experiments, we confirmed that MM-UDIPs exhibit higher cross-modal correlation between embeddings from T1 and T2 images and outperform SM-UDIPs in predicting brain-related features such as traditional IDPs. MM-UDIPs also outperform SM-UDIPs and traditional IDPs in predicting brain disorders and age. Genetic analyses of the learned embeddings revealed a substantially higher overlap of associated loci between T1 and T2 MM-UDIPs, reflecting improved alignment of shared genetic architecture across these modalities. Further protein- and drug-level analyses highlighted common targets, suggesting avenues for follow-up biological investigation.

Overall, this study presents the first multimodal contrastive learning framework for brain MRI for genetic discovery. We demonstrate that contrastive objectives can be used to learn shared representations that encode common genetic and structural properties of the human brain.

Our contributions are:

- We present, to our knowledge, the first multimodal brain IDPs, in the context of imaging genetics research.
- We provide the first direct application of momentum-encoder contrastive learning to multimodal medical images, while previous research has only applied momentum encoders to single-modal images for pretraining.
- We report a systematic biological and genetic characterization of multimodal MRI representations, while previous studies have focused mainly on performance in traditional computer vision downstream tasks, including classification, registration, and segmentation.
- We uncover loci and nominate new genes, proteins, and actionable drug targets implicated across modalities, offering translational biological hypotheses

## Results

### Overall framework of multimodal contrastive representation learning for MRI images

The overall framework is shown in **Fig 1**. The framework consists of four main components: 1) MoCoV2 contrastive pretraining on multimodal MRI images; 2) embedding extraction on an independent discovery cohort, followed by dimensionality reduction using PCA; 3) genetic analysis of the multimodal representations; and 4) follow-up analyses assessing representation alignment, predictive performance, and biological discovery potential.

**Fig 1.**
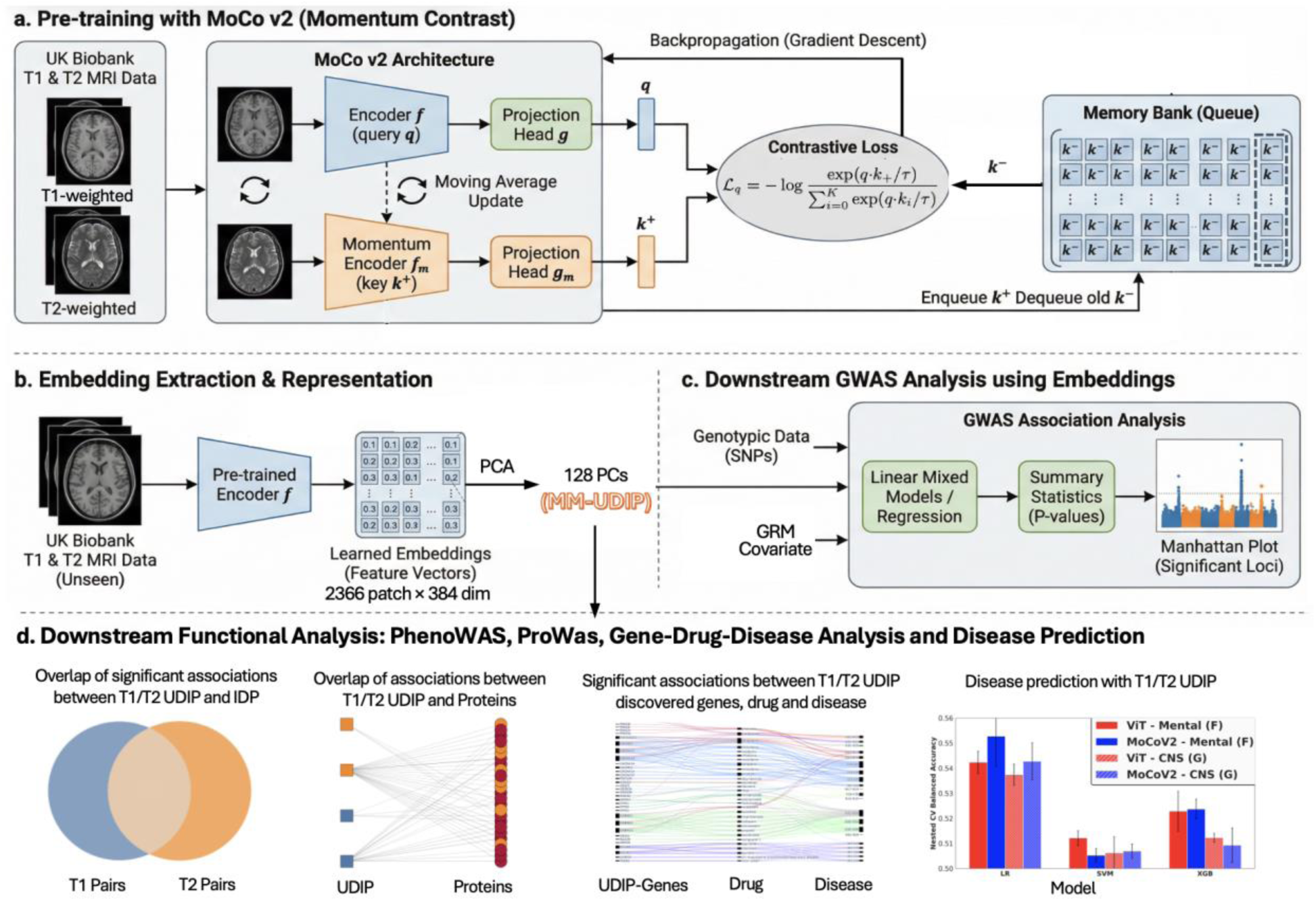
Framework of multimodal contrastive learning on MRI images. a) MoCoV2 serves as the backbone of our multimodal contrastive learning model. Here we define positive pairs as different modalities of the same individual. The framework is symmetric, allowing both T1 and T2 images to serve as the query. b) After training, the encoder *f* is applied to an independent discovery set (∼23,000 samples) to extract embeddings, followed by a dimension reduction to 128 dimensions (named MM-UDIP) using PCA for easier interpretation. c) The MM-UDIPs are analyzed using a standard genetic pipeline, performing GWAS with fastGWA and clumping loci with FUMA. d) Downstream functional analyses demonstrate that MM-UDIPs achieve superior performance in cross-modality alignment, prediction of traditional IDPs, and the identification of biologically meaningful signals.

With the goal of multimodal representation learning, contrastive learning provides a natural self-supervised framework that requires no external labels and relies solely on the intrinsic structure of the images to integrate information from multiple views of the same object. Among contrastive learning approaches, MoCoV2 offers a strong and computationally efficient baseline that remains competitive in contemporary performance benchmarks. Therefore, we adopt MoCoV2 into our contrastive learning framework and extend it to multimodal MRI images (**Fig 1a**), where paired T1- and T2-weighted MRIs from the same subject are treated as positive samples. MoCoV2 consists of two encoders: a query encoder and a key encoder, both initialized with identical weights. The key encoder is updated using a momentum-based moving average of the query encoder parameters, which stabilizes training and enables the use of a large and consistent set of negative samples.

Given an input MRI volume, the query encoder maps it to a latent representation, while the corresponding paired modality from the same subject is processed by the key encoder to form a positive pair. Negative samples are drawn from a dynamically maintained dictionary (queue) that stores representations from previous mini-batches. Contrastive learning is performed using the InfoNCE loss, which encourages the query representation to be close to its positive key while pushing it away from all negative keys in the dictionary. We construct a symmetric two-branch framework in which T1 and T2 alternately serve as the query and key. One branch treats T1 as the query and T2 as the key, while the other reverses this assignment, and the overall training objective is defined as the average loss across both branches.

In our multimodal design, T1 and T2 images from the same individual form cross-modality positive pairs, allowing the model to learn shared representation of brain structure. The queue-based contrastive mechanism enables efficient training with a large number of negative samples, improving representation quality without increasing batch size.

After training, the query encoder (*f*) was applied to an independent discovery set of 22,985 individuals to extract 384-dimensional embeddings for each of 2,366 brain patches. For each subject, the resulting 2366×384 patch-level embedding matrix was subsequently reduced to a single 128-dimensional vector using PCA (explained 73.4% variance). This dimensionality reduction was performed to reduce the computational and multiple-testing burden in downstream GWAS analyses, and to obtain a compact set of latent imaging phenotypes that facilitates downstream biological analysis. The resulting 128-dimensional representations are referred to as MM-UDIPs (**Fig 1b**). For comparison, the 128-dimensional representations learned from our previous single-modal ViT-autoencoder-based method are referred to as SM-UDIPs [14, 24]

The MM-UDIPs were then analyzed using a standard genetic analysis pipeline [14], performing GWAS with fastGWA [25] and clumping loci using FUMA [26] (**Fig 1c**). Each identified locus was extended by 125 kb on both sides, and overlaps between modalities were determined using an interval tree algorithm. Compared with the SM-UDIPs, MM-UDIPs showed a higher proportion of overlapping loci, indicating improved cross-modal genetic alignment.

Finally, we performed a series of downstream functional analyses on the MM-UDIPs (**Fig 1d**), including phenotypic correlation analysis between modalities, Phenome-wide association study (PhenoWAS), Proteome-wide association study (ProWAS), Gene-Drug-Disease Analysis, and predictions of various other phenotypes, including traditional IDPs, diseases, cognitive scores, and age. We found that MM-UDIPs have superior performance in multimodal alignment, prediction, and biological discovery.

### MM-UDIPs show alignment of genetic architecture

Here, we present multiple lines of evidence demonstrating the superior performance of the MoCoV2-based MM-UDIP over the ViT-based SM-UDIP model in aligning T1 and T2 modality embeddings, both at the feature and the genetic levels (**Fig 2a**).

**Fig 2.**
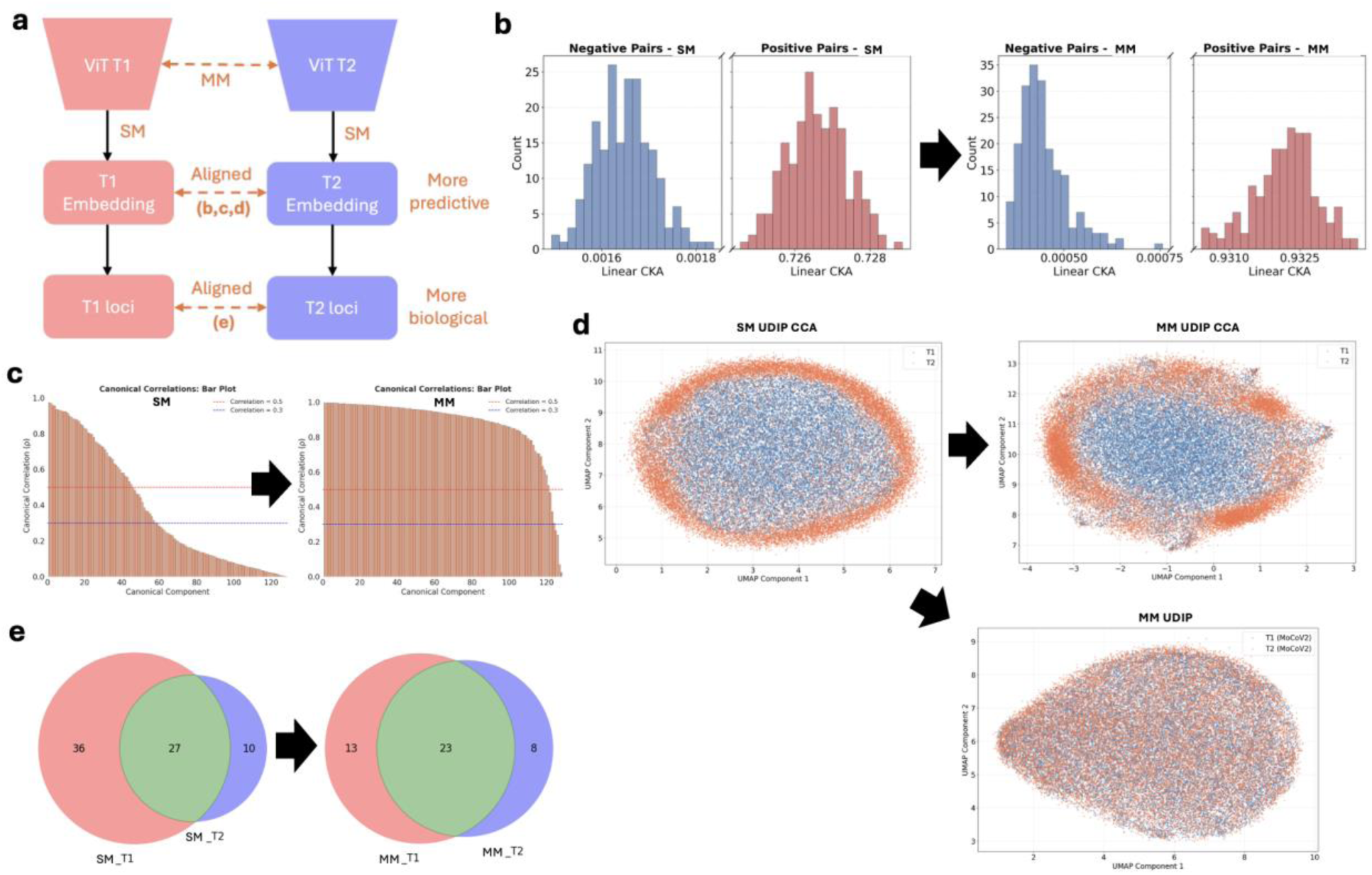
Learned multimodal embeddings present better alignment across modalities. a) MM-UDIPs demonstrate improved alignment between T1 and T2 modalities, as illustrated in panels **b-e**, and exhibit stronger predictive performance for traditional IDPs and diseases at the embedding level, as well as greater biological relevance at the genetic and loci level. For comparison, the SM model was trained using a ViT-based autoencoder with a reconstruction loss to derive 128-dimensional embeddings. b) Linear Central Kernel Alignment (CKA) was used to compare T1 and T2 UDIPs between SM and MM. Higher CKA values indicate greater similarity between the two representations. c) Canonical Correlation Analysis (CCA) was applied to evaluate T1-T2 alignment. Higher canonical correlation coefficients reflect better correspondence between embeddings. d) UMAP visualization of CCA-aligned SM-UDIPs, CCA-aligned MM-UDIPs, and original-space MM-UDIPs indicates greater T1-T2 overlap for MM-UDIPs compared with SM-UDIPs. e) Comparison of T1 and T2 loci overlap reveals that MM-UDIPs achieve a larger proportion of shared loci, indicating improved alignment between modalities at the genetic level.

Feature-level alignment is supported by two quantitative analyses (**Fig 2b-c**) and one qualitative visualization (**Fig 2d**). The quantitative measures, Central Kernel Alignment (CKA) [27] and Canonical Correlation Analysis (CCA), are widely used to assess embedding similarity. CKA is computed by taking the Frobenius inner product between the centered Gram matrices of two representations and normalizing it by the product of their Frobenius norms, so it is essentially the cosine similarity between the centered kernel matrices. CKA provides a global similarity measure that is robust to rotation and scaling, making it suitable for comparing neural network layers or embeddings [27]. CKA results show improved alignment for MM-UDIPs, with higher similarity for positive pairs (same sample, different modality: 0.727 → 0.933) and lower similarity for negative pairs (different samples, different modalities: 0.0016 → 0.0005) (**Fig 2b**). CCA analysis further confirms improved alignment of MM-UDIPs (**Fig 2c**), with larger canonical correlation coefficients across the 128 jointly embedded UDIPs between T1 and T2, as reflected by the greater area under the bars and higher mean (MM-UDIPs 0.878 vs SM-UDIPs 0.376).

Qualitative visualization using UMAP projection of T1 T2 UDIPs and CCA-transformed representations across samples further supports modality alignment (**Fig 2d**). It is important to note that SM-UDIPs derived from T1 and T2 do not reside in the same embedding space, as the ViT autoencoders for T1 and T2 were trained independently. Consequently, distances between T1 and T2 SM-UDIPs are not directly meaningful. For this reason, UMAP visualization was performed on the CCA-transformed representations, where T1 and T2 are explicitly linearly mapped into a shared coordinate space. In contrast, MM-UDIPs for T1 and T2 are produced using a shared encoder and therefore reside in a joint embedding space. Accordingly, UMAP visualizations of MM-UDIPs could be presented without requiring additional alignment. MM-UDIPs exhibit substantially greater overlap between modalities than SM-UDIPs, both before and after CCA transformation. Notably, MM-UDIPs demonstrate strong cross-modality alignment even prior to CCA, indicating that the shared encoder inherently learns modality-invariant representations.

At the genetic level, MM-UDIPs also demonstrate improved cross-modality alignment. Using a standard genetic pipeline, we identified 23 shared loci between jointly embedded T1 and T2 MM-UDIPs out of 44 total loci, representing 52.3% overlap. In comparison, SM-UDIPs shared 27 loci out of 73 total loci, corresponding to 36.9% overlap. The smaller number of loci identified for MM-UDIPs reflects an additional quality-control step introduced to mitigate inflation driven by extreme values. We observed that a small number of outliers influenced association statistics (**Supp Fig 2-4**). Specifically, removing outlier sample values exceeding 5 standard deviations reduced the number of T1 loci from 219 to 36, highlighting the necessity of this quality control step.

### Phenotypic annotation of MM-UDIPs using conventional imaging traits

The primary goal of conducting a PhenoWAS is to interpret UDIPs and reveal their biological significance. To achieve it, we performed a comprehensive phenome-wide association analysis between UDIPs and traditional IDPs for both MM-UDIPs and their reconstruction-based predecessor, SM-UDIPs. First, we examined the overall distribution of association p-values between UDIPs and IDPs across modalities (T1 and T2) and models (SM and MM), with covariates corrected (**Fig 3a**). After Bonferroni correction (P < 0.05/128/179), we identified 9,053 and 9,044 significant UDIP-IDP associations for MM T1 and T2, respectively, compared with 13,754 and 12,303 for SM T1 and T2. Although the number of significant associations differs between MM and SM, the association strength (p-values) for a given traditional IDP is largely comparable across the two models.

**Fig 3.**
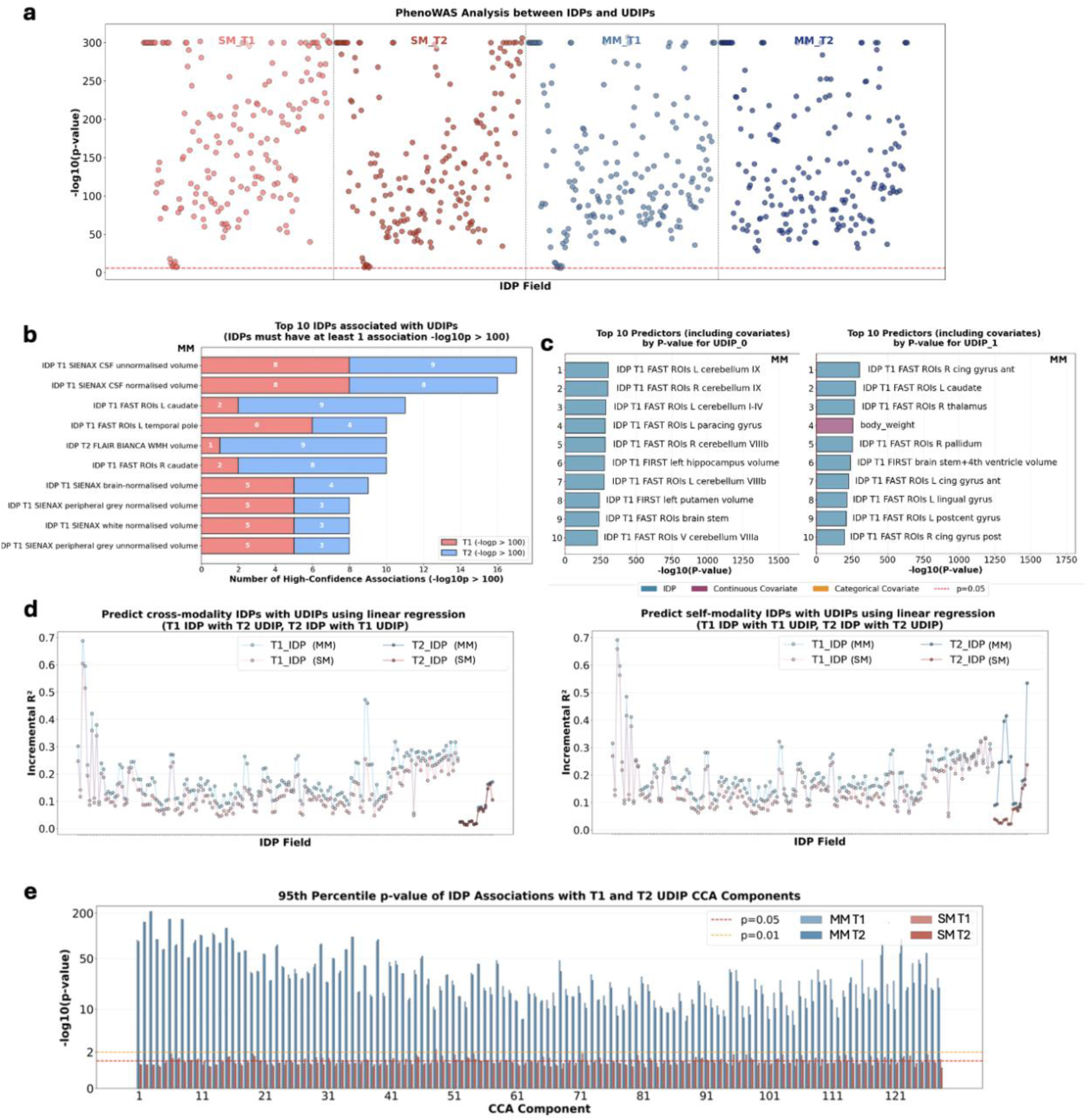
Phenotype analysis. a) Scatter plot of −log _10_ *p*-values from univariate analyses between traditional IDPs and UDIPs, accounting for covariates. The red line indicates the Bonferroni correction p-value of 0.05/128/179. b) traditional IDPs ranked by the number of associations exceeding −log _10_ *p* > 100. c) Associations of UDIPs with covariates and traditional IDPs calculated separately, showing the top 10 significant associations for the first two UDIPs (UDIP_0 and UDIP_1). d) Comparison of predictive performance shows that MM-UDIPs achieve greater improvements in traditional IDP prediction-both within the same modality and across modalities-relative to SM-UDIPs. e) Comparison of 95^th^ percentile p-value of traditional IDP association with CCA components from UDIPs.

Next, we examined the most significant UDIP-IDP associations (p < 10⁻¹⁰⁰) and found that MM-UDIPs align more closely with traditional volume-based IDPs (**Fig 3b**). Top associations for MM-UDIPs included T1 CSF volume (both normalized and unnormalized), T1 left and right caudate, and T1 left temporal pole. Notably, the caudate and temporal regions are adjacent to CSF, reflecting a consistent biological pattern.

To further assess whether UDIPs capture imaging-specific information rather than confounding factors, we calculated associations between UDIPs and covariates (**Fig 3c**). For example, the first two T1 MM-UDIPs, which explain the 18.2% and 6.3% variance in T1 data, show their strongest associations with imaging features rather than covariates such as age. This trend can be further explored across additional top UDIPs (**Supp Fig 5**)

Importantly, we assessed the predictive power of MM-UDIPs for both same-modality and cross-modality traditional IDPs, compared with SM-UDIPs, using incremental *R*^2^ from multivariable prediction models (**Fig 3d**). The incremental *R*^2^ quantifies the additional variance explained by UDIPs beyond covariates alone. We found that MM-UDIPs have higher mean incremental *R*^2^ in all four groups of predicting both self-modality and cross-modality traditional IDPs (T1_IDP_T1_UDIP: MM 0.19, SM 0.16; T1_IDP_T2_UDIP: MM 0.18, SM 0.14; T2_IDP_T1_UDIP: MM 0.07, SM 0.06; T2_IDP_T2_UDIP: MM 0.22, SM 0.07). We also applied partial F-tests (omnibus tests) to compare the models, which showed higher statistics for MM-UDIPs in predicting both self-modality and cross-modality traditional IDPs (**Supp Fig 6**). This indicates that the coefficients of MM-UDIPs are less likely to be zero. Together, these results demonstrate that MM-UDIPs provide stronger predictive performance for traditional IDPs than SM-UDIPs.

Finally, we sought to demonstrate that MM learns shared, biologically meaningful latent structure across imaging modalities. Our central hypothesis was that MM-UDIPs would exhibit strong canonical correlations between T1 and T2, and that the resulting CCA components would map consistently onto traditional IDPs. This rationale follows from canonical correlation analysis, which addresses a core question in multimodal representation learning: which linear combinations of T1 features covary maximally with linear combinations of T2 features across individuals. Each CCA mode captures a latent source of inter-subject variation that is jointly expressed in both T1 and T2 embeddings. When these shared embedding dimensions show strong associations with known traditional IDPs, they indicate that the learned representations encode meaningful anatomical variation.

To test the hypothesis, we performed univariate association analyses between CCA components of UDIPs and traditional IDPs, and compared MM with SM using the 95th-percentile association p-values (**Fig 3e**). MM exhibited substantially stronger associations than SM. As expected, association strength within MM decreased slightly with increasing CCA component index, reflecting diminishing shared variance. In contrast, SM associations remain stable near nominal significance thresholds of 0.05 to 0.01.

Taken together, these findings demonstrate that MM captures more robust and biologically meaningful shared latent structure between T1 and T2 modalities than reconstruction-based SM.

### Proteomic profile of the MM-UDIPs

To further characterize the shared biological signal captured by MM-UDIPs, we conducted a ProWAS, examining associations between 2,457 Olink plasma proteins and each of the 128-dimensional MM-UDIPs (Methods).

After Bonferroni correction (P < 0.05/128/2,457), we identified 19 and 27 significant protein associations for jointly embedded T1 and T2 MM-UDIPs, respectively (**Fig 4a**). Among these, 17 proteins were shared between T1 and T2, out of a total of 25 significant proteins, while 8 proteins were unique to T2 and none were unique to T1 (**Fig 4a**).

**Fig 4.**
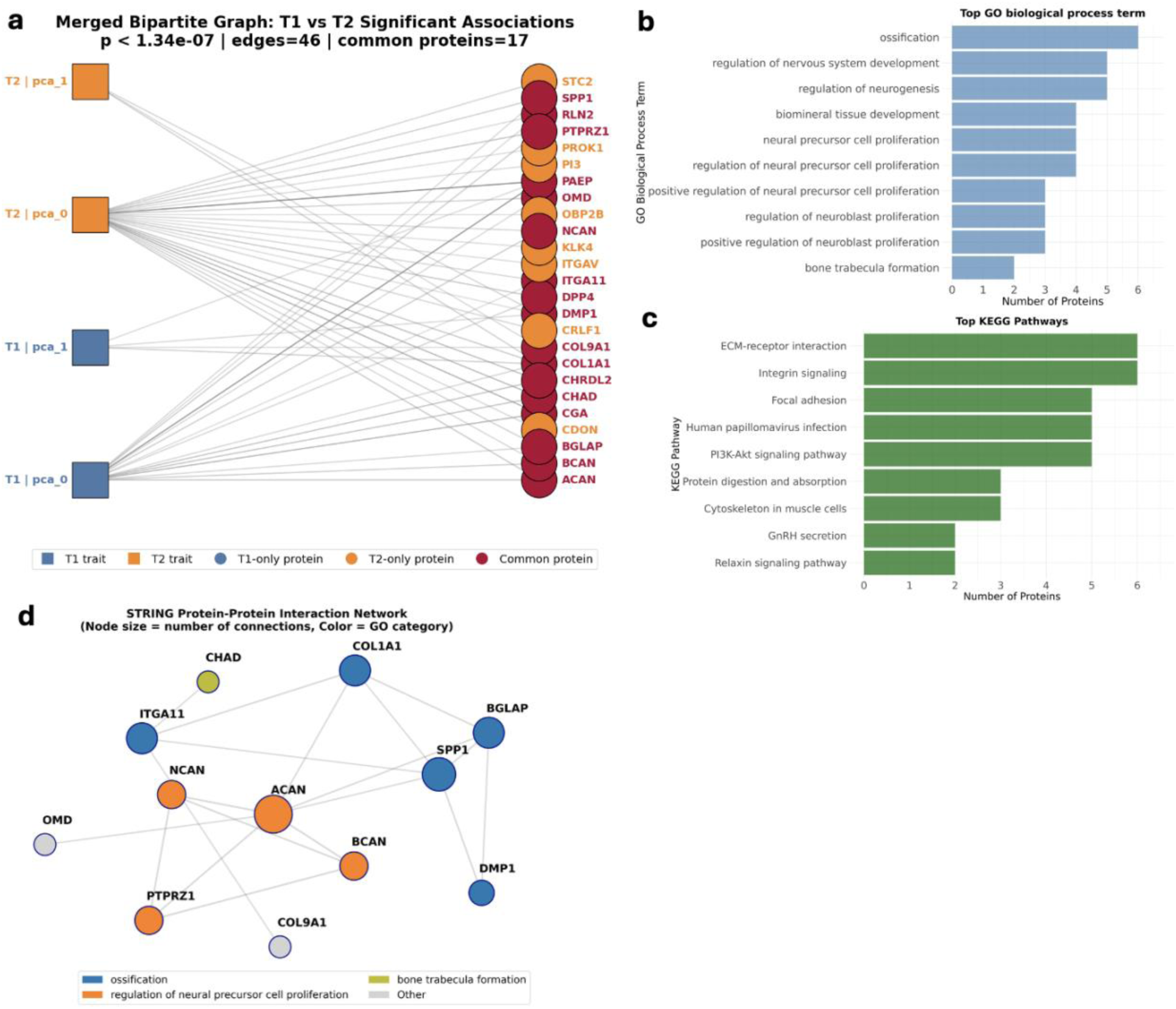
Proteomic analysis on MM-UDIPs. a) Overlap of significantly associated proteins between jointly embedded T1 and T2 MM-UDIPs. Line color indicates association significance, with darker lines representing lower p-values. Only the first two UDIPs, corresponding to the first two principal components, show significant protein associations. b) Top GO pathways ranked by the number of associated proteins. c) Top KEGG pathways ranked by the number of associated proteins. Both the top GO and KEGG pathways are related to neural system development. d) STRING protein interaction network illustrates the interactions among the identified common proteins, with colors indicating their associated GO pathways.

Using the 17 proteins shared between jointly embedded T1 and T2 MM-UDIPs, we performed pathway enrichment analysis based on the GO and KEGG databases. GO pathway names tend to be more descriptive biologically, and the top GO pathways are primarily related to neural cell development (**Fig 4b**). The top-ranked GO pathway, ossification, refers to the formation of new bone by osteoblasts, which aligns with bone-related loci and genes, such as **BMP4**, identified in later genetic analyses.

KEGG pathway names are more molecularly oriented, and the top 10 KEGG pathways similarly highlight neural system development (**Fig 4c**). A more detailed discussion incorporating newly discovered genes is provided in the subsequent genetic analysis section.

The STRING protein interaction network further illustrates relationships among the identified proteins (**Fig 4d**). Notably, **ACAN**, **BCAN**, and **NCAN** are key brain-specific extracellular matrix (ECM) components that encode chondroitin sulfate proteoglycans. These molecules form the core of perineuronal nets, specialized ECM structures that stabilize synapses and regulate plasticity in the mature human cortex [28]. These proteins closely correspond to the gene list identified in our genetic analysis, as described in the following section.

### Genetic analysis of MM-UDIPs

Most importantly, we explored the genetic architecture of MM-UDIPs using a standard genetic analysis pipeline (Methods). Our goal was to characterize the novel loci and genes identified by the combined T1/T2 representation, which captures more comprehensive information about brain structure, making the discovered loci potentially more robust and biologically meaningful.

Among the loci identified using MM-UDIPs, 23 loci are shared between T1 and T2 out of a total of 44 loci, representing 52.3% overlap. By comparison, SM-UDIPs share 27 loci out of 73 total loci, corresponding to 36.9% overlap. The distribution of MM T1 and T2 loci is shown in **Fig 5a**. Annotation using the GWAS Catalog revealed that, for the 44 loci identified with MM-UDIP, 55.2% of the top five traits per locus (ranked by p-value) are related to brain or vertex phenotypes. In contrast, 52.1% of traits associated with SM T1 and T2 loci are brain- or vertex-related. These results suggest that loci discovered with MM-UDIP are slightly more biologically meaningful. It should be noted that some loci identified by MM-UDIP are novel and never been reported, so simply calculating the percentage of loci containing GWAS Catalog entries with brain-related traits may be misleading. Using the proportion of the top five GWAS Catalog traits within each locus provides a more accurate assessment of biological relevance.

**Fig 5.**
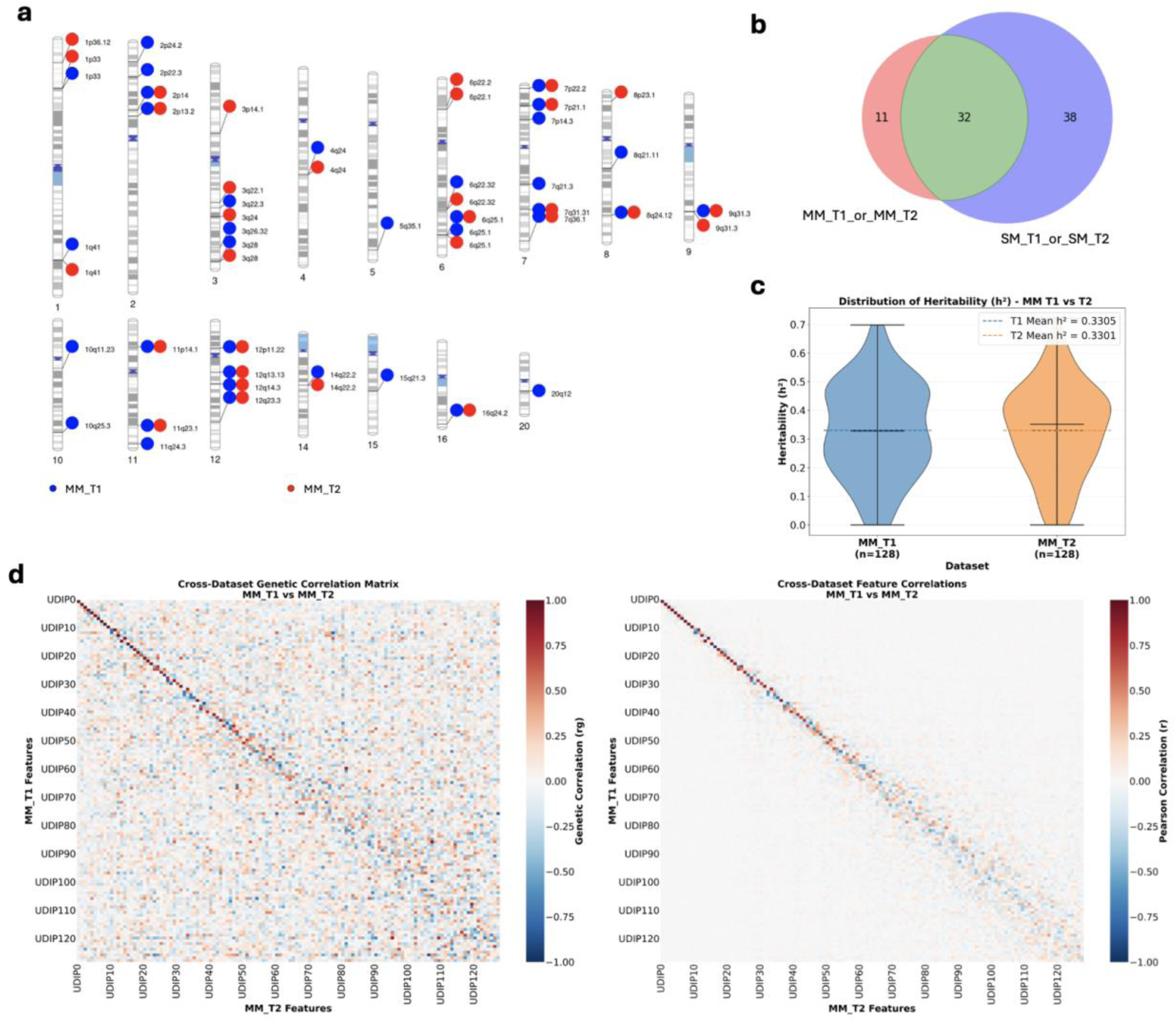
Genetic analysis on UDIPs. a) Genomic loci jointly and separately associated with T1 and T2 MM-UDIPs. Significant loci were determined using Bonferroni correction (two-sided P < 5 × 10⁻⁸/128). b) Newly identified loci that were not reported in the previous ViT-based SM analysis. c) SNP-based heritability (ℎ^2^) estimates for the 128 jointly embedded T1 and T2 MM-UDIPs, calculated using GCTA software. d) Genetic correlation between the 128 T1 MM-UDIPs and 128 T2 MM-UDIPs.

We identified 11 novel loci with MM-UDIP that were not reported by the previous SM ViT model on T1 and T2 (**Fig 5b**). Among these, 7 loci were also not reported in prior CNN-based analyses, and after cross-referencing with previously reported traditional IDP loci, 6 loci appear to be entirely novel. The detailed overlap between MM-UDIP-discovered loci and loci from previous methods is provided in **Supp Table 1**.

For the 11 novel loci, the top five previously reported GWAS Catalog traits are diverse. Some are directly related to brain structure, such as white matter microstructure, brain morphology, or subcortical volume (e.g., rs13388394). Others are associated with metabolic traits, including non-HDL cholesterol levels (rs60852193), liver enzyme alkaline phosphatase (rs10783562), or platelet-to-lymphocyte ratio (rs238871).

More interestingly, the gene list derived from the genetic analysis exhibits substantial functional convergence with the proteins identified as significant in the ProWAS, despite a lack of direct gene overlap. This convergence is especially evident in pathways related to the brain extracellular matrix (ECM) and proteoglycan biology. The protein list is strongly enriched for ECM and brain-specific ECM components (ACAN, BCAN, NCAN), which encode chondroitin sulfate proteoglycans that are core constituents of perineuronal nets-specialized ECM structures that stabilize synapses and regulate plasticity in the mature human cortex [28]. In addition, the protein list includes PTPRZ1, a receptor tyrosine phosphatase abundantly expressed in the developing human cerebellum and implicated in oligodendrocyte development and neural communication [29]. In contrast, genes from the genetic analysis predominantly implicate regulatory mechanisms of ECM organization rather than structural ECM components. These include ADAMTS8 [30], an ECM-degrading protease; BMP4, a morphogen linked to ECM-mediated signaling [31, 32]; and NUAK1, which modulates cell adhesion, energy metabolism, and polarity [33, 34]. These lines of evidence point to shared involvement in neurodevelopmental processes related to ECM regulation, cell adhesion, and signaling.

Beyond ECM-related processes, GWAS and ProWAS discovered gene sets also converge on neurodevelopmental signaling pathways. The genetic analysis highlights genes such as BMP4, ALDH1A2, WNT16, LPAR1, and NUAK1, which are central to developmental patterning, cortical development, and axon guidance. Correspondingly, the protein list includes CHRDL2, a known antagonist of BMP signaling [35], as well as SPP1 (osteopontin), which participates in cell signaling and neuroinflammatory processes [36], and RLN2, which is conceptually related to Reelin-family signaling. Collectively, these findings indicate shared involvement in BMP, WNT, and ECM-related developmental pathways, suggesting coordinated regulation of brain structure and maturation across genetic and proteomic levels.

We calculated SNP-based heritability to quantify the proportion of phenotypic variance explained by common genetic variants across the genome(**Fig 5c**). Among the 128 MM-UDIPs, GCTA [37] identified significant heritability in 29 T1 UDIPs and 20 T2 UDIPs (T1/T2 ℎ^2^ = 0.33 ± 0.16; P < 0.05/128). The h2 statistics cannot be directly compared with previous model and traditional IDPs as they are highly correlated, whereas MM-UDIP is decorrelated by PCA. The heritability estimates reported from MM-UDIP more accurately reflect true genetic contributions rather than potential inflation caused by correlated features. We further conducted an ablation study in which the same procedure used for MM-UDIPs, PCA rather than average pooling, was applied to generate 128 ViT-based SM-UDIPs, enabling direct comparison with MM-UDIPs. This PCA-based SM-UDIP achieved mean heritability estimates of 0.11 for T1 and 0.09 for T2 (**Supp Fig 7**), substantially lower than those of MM-UDIPs, which reach 0.33 for both T1 and T2. To further characterize the genetic architecture of MM-UDIPs from T1 and T2 modalities, we estimated the genetic correlation between the two sets of UDIPs (**Fig 5d**), as well as showing the phenotype correlation between T1 and T2 MM-UDIPs. We observed that corresponding leading UDIPs from T1 and T2 showed stronger genetic correlations. Notably, the mean absolute genetic correlation for the corresponding T1–T2 UDIPs (diagonal entries in the heatmap) is greater than that of the phenotype itself (0.37 vs. 0.34), indicating that the alignment between T1- and T2-based UDIPs extends beyond the feature level to the genetic level. This finding also suggests that loci identified from MM-UDIP are likely to be biologically meaningful, as they derive from embeddings that are strongly correlated at the genetic level.

### MM-UDIPs Drug-target annotation and prediction for systemic diseases, cognition and age

To explore the translational applications of the MM-UDIPs, we performed a gene-drug-disease enrichment analysis [38] using Multi-marker Analysis of GenoMic Annotation (MAGMA) genes (p < 0.05/18,943 for 18,943 genes) associated with the 128 MM-UDIPs to queue the related drugs and diseases from the DrugBank database [39] and Therapeutic Target Database [40]. This analysis generated a gene-drug-disease network to identify potentially repurposable drugs, a strategy that has been shown to improve success rates in drug development [41, 42].

The gene-drug-disease network identified 989 significant pairs of gene-drug-disease interactions (p<0.05) between 80 genes, 367 drugs and 28 International Classification of Diseases (ICD) disease categories (**Fig 6a**) for T1 and 1,001 significant pairs between 75 genes, 370 drugs and 32 ICD diseases for T2 (**Fig 6b**). Here we plot the top 30 drugs with maximum numbers of associations and mark the common drugs between T1 and T2 analysis in red. The following are the illustrations of several research regard those common drugs and/or small molecules. For example, the GABRB gene family served as the target gene for mental and neural disorder F30-39, G40-47, G60-64, G90-99, which could be treated with ‘topiramate’, ‘iorazepam’, ‘pentobarbital’,‘meprobamate’,‘butalbital’. The GABRB1, GABRB2, and GABRB3 genes are central to assembling functional GABA_A receptors. Drugs like topiramate, lorazepam, pentobarbital, meprobamate, and butalbital all modulate GABA_A receptor activity, enhancing inhibitory neurotransmission. Through this shared mechanism, they can treat conditions involving neuronal hyperexcitability and dysregulated inhibitory signaling, such as certain epilepsies, anxiety disorders, and other mental/neural disorders.

**Fig 6.**
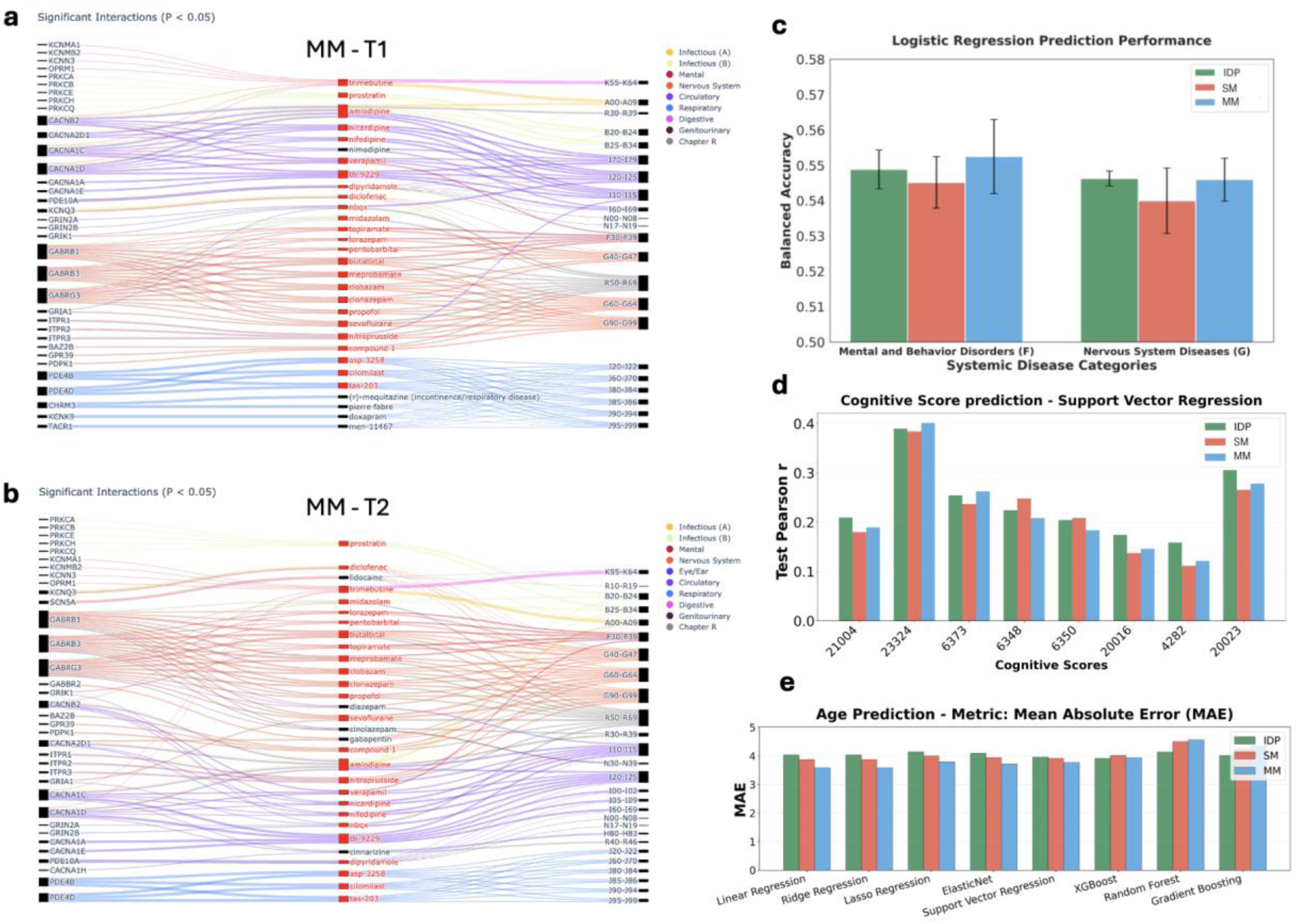
The translational application potential of MM-UDIP. a) Gene-drug-disease enrichment analysis [38] using MAGMA genes associated with the 128 T1 MM-UDIPs. Red lines indicate associations with brain-related disorder with ICD codes F and G. b) Gene-drug-disease enrichment analysis using MAGMA genes associated with the 128 T2 MM-UDIPs. c) Comparison of disease prediction performance using traditional IDPs, SM-UDIPs, and MM-UDIPs, evaluated by nested cross-validation balanced accuracy. d) Comparison of cognitive score prediction performance using traditional IDPs, SM-UDIPs, and MM-UDIPs, evaluated by Pearson r on the test set. 8 cognitive scores include Matrix Pattern Completion (Field ID: 21004); Digit Symbol Substitution Test (DSST) (Field ID: 23324); Tower Rearranging (Field ID: 6373); Trail Making Test A (TMT-1) (Field ID: 6348); Trail Making Test B (TMT-2) (Field ID: 6350); Fluid Intelligence (Field ID: 20016); Numeric Memory (Field ID: 4282); Reaction Time (Field ID: 20023). e) Comparison of age prediction performance using traditional IDPs, SM-UDIPs, and MM-UDIPs, evaluated by mean absolute error.

We also noticed that there is a drug named with ‘compound 1’ that is associated with neural disease G60-64 and G90-99 and associated with gene BAZ2B, GPR39, PDPK1. ‘Compound 1’ is a placeholder name in Therapeutic Target Database (TTD) for research compounds without official names. There are three entries D0A4RF, D0H3UW, and D0FE7H all refer to “compound 1” variants linked to neurological conditions: D0A4RF (PubChem CID 657061) is indicated for pain, D0H3UW (CID 44409861) is reported for Alzheimer’s disease, and D0FE7H (CID 9876120) is listed for multiple sclerosis. Notably, the pain indication falls under ICD neurology categories (e.g. G60-G64/G90-G99, neuropathies and nervous system disorders). These TTD entries suggest a potential link between these research compounds and neural disorders.

We evaluated the predictive power of MM-UDIPs relative to SM-UDIPs and traditional IDPs across multiple tasks, including two systemic disease categories (based on ICD-10 codes), eight cognitive scores, and age (**Fig 6c-e**). Case and control group definitions are provided in the Methods. These analyses test the hypothesis that multimodal contrastive learning produces embeddings that are better aligned and more predictive than those derived from a single modality. Overall performance was modest, reflecting the inherent difficulty of the tasks and class imbalance. Proteomics data were excluded due to high rates of missing values across individuals and proteins.

We first assessed the predictive power of multimodal imaging features by combining jointly embedded T1 and T2 MM-UDIPs for classification of two disease categories (**Fig 6c, Methods**). Under nested cross-validated logistic regression, MM-UDIPs outperformed SM-UDIPs in predicting both mental and behavioral disorders (ICD: F) and nervous system diseases (ICD: G). Compared with traditional IDP features, MM-UDIPs showed superior performance for mental and behavioral disorders and comparable performance for nervous system diseases (Mean Balanced Accuracy, F: MM-UDIP 0.553, SM-UDIP 0.542; G: MM-UDIP 0.543, SM-UDIP 0.537). Among all models tested, including logistic regression, SVM, and XGBoost, logistic regression achieved the highest accuracy (**Supp Fig 8**).

Next, we evaluated the predictive power of MM-UDIPs for eight cognitive scores (**Methods**), comparing them with SM-UDIPs and traditional IDPs (**Fig 6d**). We report test performance using support vector regression, which outperformed linear regression with regularization. Across the eight cognitive scores, MM-UDIPs outperformed SM-UDIPs in six cases and exceeded traditional IDP performance in two. The strong performance of traditional IDPs likely reflects their higher feature dimensionality compared with the 128 features used for UDIPs.

Finally, we evaluated age prediction performance using MM-UDIPs across multiple models, compared with SM-UDIPs and traditional IDPs (**Fig 6e**). AI-derived brain age has emerged as a valuable biomarker for assessing brain health [43–45], and several studies highlight the use of the biological age gap (BAG) due to its rich information spanning multiple modalities, organs, and omics [46–49]. Across eight models, MM-UDIPs achieved lower mean absolute error in six, outperforming both SM-UDIPs and traditional IDPs.

## Discussion

In this paper, we demonstrate that MoCoV2 effectively generates multimodal-aligned UDIPs (MM-UDIP) that substantially outperform single-modal UDIPs (SM-UDIP) across representation alignment, predictive utility, and biological discovery (**Table 1**). MM-UDIP exhibits markedly stronger cross-modal consistency, evidenced by higher positive-pair similarity and lower negative-pair similarity in CKA (P = 0.932 vs. 0.726; N = 0.0004 vs. 0.0016), a much higher mean CCA correlation (0.878 vs. 0.376), and clearer structural agreement in low-dimensional embeddings (high vs. low UMAP overlap). These alignment gains translate into downstream benefits: MM-UDIP achieves superior performance in traditional IDP prediction, disease classification, cognitive score prediction (winning in 6/8 tasks), and age estimation, consistently exceeding SM-UDIP across all evaluated metrics.

**Table 1.**
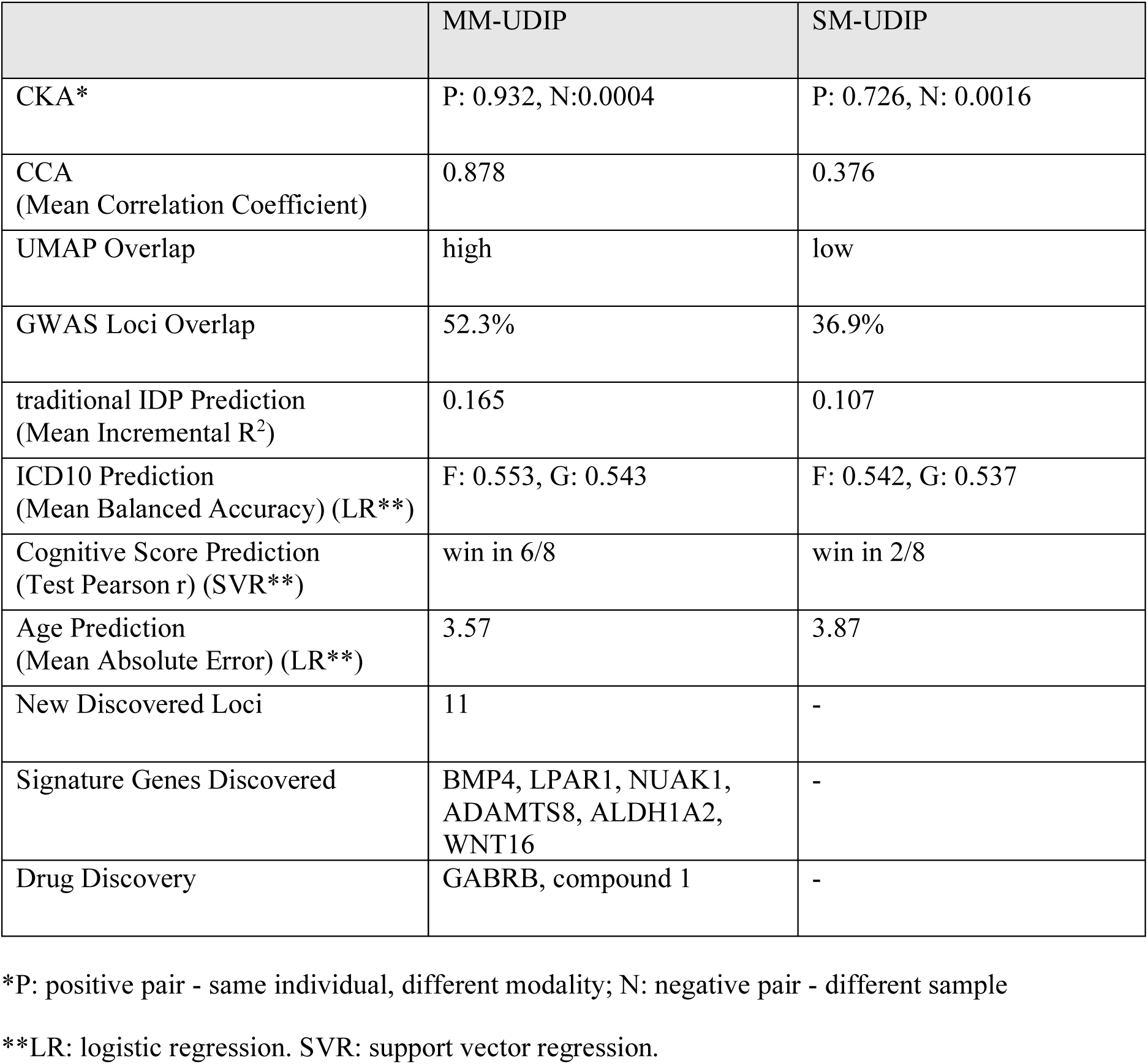
Comparison of performance between MM-UDIP and SM-UDIP.

Crucially, the advantages of MM-UDIP extend beyond prediction to biological and translational discovery, which is largely absent in the single-modality setting. MM-UDIP recovers a substantially higher overlap with known GWAS loci (52.3% vs. 36.9%) and uniquely enables the identification of 11 novel loci, along with biologically interpretable signature genes (e.g., BMP4, LPAR1, NUAK1, ADAMTS8, ALDH1A2, WNT16) and candidate drug targets (GABRB, compound 1). These findings indicate that multimodal alignment does not merely improve numerical performance but also enhances genetic signal coherence, allowing MM-UDIP to capture shared biological mechanisms across imaging modalities that remain inaccessible to SM-UDIP. Together, these results establish MM-UDIP as a more expressive, biologically grounded representation that advances both predictive modeling and discovery in imaging genetics.

While these results are promising, there remains substantial room for further improvement. First, we describe the model design and the rationale behind key parameter choices. Notably, the MoCoV2 framework adopted in this study differs from the original MoCoV2 formulation, in which multiple augmented views of the same image are treated as positive pairs. Instead, we define multimodal views of the same subject (i.e., paired T1- and T2-weighted MRI) as positive pairs, while all other combinations,including different subjects and different modalities,are treated as negative pairs. To the best of our knowledge, this represents the first application of directly learning multimodal image representations using a momentum encoder. This contrasts with prior approaches that pretrain momentum encoders on single modalities and subsequently concatenate the learned embeddings, which have been referred to as multimodal momentum learning [50, 51].

The choice of MoCoV2 over other self-supervised learning like SimCLR [52, 53] or DINO [54–56] is primarily motivated by computational efficiency. MoCoV2 maintains a dictionary-based negative queue, which effectively enlarges the batch size while substantially reducing GPU memory requirements. This advantage is particularly important for high-resolution 3D MRI data, where memory constraints limit the number of samples per batch.

PCA is employed as a dimensionality reduction step to obtain 128-dimensional subject-level embeddings from the patch-level representations of size 2366×384. PCA provides a simple and effective way to compress high-dimensional features. Another straightforward option is to perform average pooling directly on the 384-dimensional patch embeddings. Although this approach avoids dimensionality reduction altogether, it results in a higher-dimensional subject-level representation, which increases the complexity of downstream analysis and reduces interpretability. We also tried reduce the training dimension to 128 followed by an average pooling across the 2,366 patches, thereby eliminating the need for PCA. However, in our experiments, models trained with a reduced latent dimensionality of 128 exhibited inferior performance (**Supp Note 1**), suggesting that higher-dimensional patch representations (384 dimensions) are necessary to preserve sufficiently rich and discriminative information.

Consequently, using PCA to project the pooled representations into a 128-dimensional space represents a pragmatic compromise. This strategy retains much of the informative variance captured by the higher-dimensional embeddings while producing a more compact and interpretable representation that is better suited for subsequent analysis and comparison.

Second, there are some future improvement on the model architecture, inspired by other categories of self-supervised learning (SSL). SSL has become a widely adopted approach for learning feature representations from large-scale unlabeled data. Existing SSL methods can be broadly grouped according to their pretext tasks or loss functions. First, generative SSL methods rely on autoencoder-style architectures to reconstruct part or all of the input data, with learning driven by reconstruction error.

Second, discriminative SSL methods train models to solve predefined prediction tasks, such as recovering the spatial arrangement or ordering of image patches. Third, contrastive learning-based SSL methods encourage consistency between different augmented views of the same sample while separating representations of different samples in the embedding space. Fourth, adversarial SSL methods incorporate adversarial objectives to remove nuisance or unwanted information from learned representations while preserving predictive structure under self-supervised supervision. Among these categories, generative and contrastive approaches have been the most influential and have driven substantial progress in representation learning. Prominent examples include SimCLR [52, 53], MoCo [23, 57, 58], BYOL [59], SimSiam [60], DINO [54–56], and VICReg [61]. All these SSL methods have the potential to be applied for generating multimodal brain MRI representations.

In fact, SSL has already been applied in several contexts within MRI analysis. For the generative-based SSL, our lab has previously applied an autoencoder-based method for the reconstruction of single modality MRI T1 or T2 images and subsequently leveraged these representations for genetic association analyses [14, 24]. For the contrastive-learning-based SSL, multimodal MRI research encompasses a range of task categories that leverage complementary information across imaging contrasts for different objectives. One major category is MRI synthesis and imputation, which aims to reconstruct missing or enhance existing contrasts by modeling cross modality dependencies, commonly using generative adversarial networks, hybrid fusion architectures, or transformer-based models to improve reconstruction fidelity and data completeness [62–66]. Closely related to synthesis, cross modality translation focuses on learning mappings between MRI contrasts or domains, often under incomplete input settings, to enable modality conversion or multi domain completion [67, 68]. Another important category combines multimodal fusion and prediction, where multiple MRI sequences are jointly integrated to learn shared or complementary representations for downstream tasks such as disease classification, prognosis, or consistency prediction, either through multimodal pretraining or radiomics based feature aggregation to improve predictive performance and robustness [69–71]. In addition, multimodal MRI is widely used in segmentation and registration tasks, where information from multiple contrasts improves anatomical delineation and spatial alignment through interactive feature learning or contrastive representation learning [72, 73]. Across these task categories, most existing approaches primarily emphasize image level reconstruction quality or task specific performance by exploiting correlated targets across modalities, with limited attention to the biological interpretation of modality interactions or the physiological meaning of the learned multimodal representations.

The model employed in this work is a purely encoder-based framework that focuses on learning discriminative latent representations without explicitly constraining their ability to reconstruct the input data. While this design is effective for contrastive representation learning, it may overlook complementary information captured by generative or reconstruction-based objectives. Incorporating a reconstruction component-such as a decoder trained to recover the original input images-could provide additional structural regularization on the learned embeddings.

Specifically, integrating a reconstruction loss would encourage the latent representations to preserve fine-grained anatomical and intensity information, which is particularly important for high-dimensional 3D neuroimaging data. This hybrid formulation would combine the strengths of contrastive learning, which promotes invariance and discriminability, with reconstruction-based learning, which enforces fidelity to the underlying data distribution.

Such an extension naturally aligns with our previous publication [14], where reconstruction objectives were shown to capture biologically meaningful brain structure. A unified framework that merges the encoder-decoder architecture from our prior work with the contrastive learning strategy presented here could therefore yield more informative and stable representations. Moreover, this combined approach may improve generalization, interpretability, and robustness to noise, especially in settings with limited labeled data.

Discriminative-based SSL focuses on learning embeddings by solving tasks that require distinguishing between different types of input transformations or configurations. For example, one could train the model to predict the correct order of shuffled image patches, forcing the network to capture meaningful spatial and structural relationships within the brain. Such tasks encourage the latent representations to encode rich contextual and anatomical information beyond what standard contrastive learning achieves.

Adversarial SSL, on the other hand, introduces a generative adversarial component, such as a GAN, to regularize the learned embeddings. By training the encoder to produce features that are difficult for a discriminator to distinguish from real data, the model can learn more robust and realistic representations that better capture the underlying data distribution.

Beyond these, additional complementary objectives-such as segmentation loss or perceptual loss-could be incorporated. Segmentation loss would enforce anatomical consistency and structural fidelity, guiding the embeddings to reflect meaningful tissue or region-level information. Perceptual loss, often computed using features from a pretrained network, encourages the model to retain high-level structural and textural characteristics of the input images.

Exploring these alternative SSL strategies could provide richer, more biologically relevant representations, improve cross-modal alignment, and enhance downstream tasks such as phenotype prediction, disease classification, or genetic association analysis.

Third, the proposed framework could be further extended to incorporate all four or five major MRI modalities available in large-scale neuroimaging datasets. In this study, we focus on T1- and T2-weighted structural MRI, as these modalities are the most widely studied and provide clear and well-interpreted anatomical information, making them a natural starting point for representation learning and cross-modal alignment. However, additional modalities such as Functional MRI (fMRI) and diffusion tensor imaging (DTI) remain unexplored in the current work. These modalities capture complementary aspects of brain organization that are not directly observable from structural images alone. For example, fMRI measures brain activity indirectly through blood-oxygen-level-dependent signals, enabling the mapping of task-related and resting-state functional dynamics, while DTI characterize the diffusion of water molecules to infer white-matter microstructure and structural connectivity. Incorporating these modalities could substantially enrich the learned representations by integrating anatomical, functional, and connectivity-based information. Diffusion MRI can complement both structural and functional findings by revealing whether abnormalities across distributed regions are supported by disruptions in underlying white-matter connectivity. Similarly, MRS can add metabolic and cellular information that helps interpret volumetric or functional differences in terms of neuronal density, integrity, or biochemical status. Through such cross-modal complementarity, multimodal MRI enables a more complete and biologically grounded interpretation of observations derived from any single modality. Quantitative MRI sequences further enhance this approach by augmenting conventional structural MRI with biophysically interpretable tissue parameters, improving characterization of inter-regional heterogeneity and inter-individual variability in the human brain [74–76].

Extending the model to a multimodal setting also introduces new methodological challenges, including differences in spatial resolution, temporal structure, noise characteristics, and alignment across modalities. Addressing these challenges would require careful architectural design and modality-specific encoders, as well as strategies to enforce coherent cross-modal representation learning. Nonetheless, such an extension holds promise for learning more comprehensive and biologically meaningful brain representations that better reflect the multifaceted nature of brain organization.

Finally, beyond the predictive gains introduced by MM-UDIPs and the potential methodological improvements, their most compelling contributions lie in biological discovery. Among the 11 loci uniquely identified by MM-UDIP, we highlight three genes of particular interest due to their established roles in neural development, brain organization, and disease-related pathways, offering biologically interpretable insights and motivating further downstream analyses by researchers from diverse biological backgrounds.

LPAR1 (Lysophosphatidic Acid Receptor 1, Edg2), for example, encodes the lysophosphatidic acid (LPA) receptor 1, a GPCR for the lipid signaling molecule LPA. LPAR1 is highly expressed in the brain [77]. It is found abundantly in the cortical ventricular zone during development and in white matter, and is present on neurons, astrocytes, oligodendrocytes and microglia. LPA-LPAR1 signaling has been shown to regulate neural progenitor cell proliferation, neuronal migration, axon guidance, and myelination. In human studies and animal models, LPAR1 activity influences cortical development and is implicated in psychiatric conditions. For example, excessive LPAR1 signaling can lead to developmental brain abnormalities. Thus, LPAR1’s primary role is neural: it mediates LPA-induced neuronal differentiation and glial function in the CNS [77].

NUAK1 (also ARK5) is an AMPK-related kinase implicated in neuronal development. Human genetic evidence supports its connection to neurodevelopmental disorders: rare de novo variants in NUAK1 have been identified in individuals with autism spectrum disorder (ASD) in large sequencing cohorts, and NUAK1 is scored as a strong candidate risk gene in curated ASD gene databases based on these findings and mouse functional data linking variant effects with neural phenotypes. Functional studies in neuronal systems show that NUAK1 controls cortical axon branching by locally modulating mitochondrial metabolic functions, effectively linking energy homeostasis and neuronal morphogenesis [34]. Mouse models demonstrate that Nuak1 haploinsufficiency impairs cortical connectivity and leads to behavioral abnormalities relevant to social novelty and cognitive function, consistent with human ASD associations [78]. While NUAK1 also plays roles in non-neural processes such as cell adhesion, stress responses, and cancer biology, its most striking functions appear neuronal, orchestrating energy-dependent axon branching and polarity establishment during cortical development.

BMP4 (Bone Morphogenetic Protein 4) is a secreted member of the TGF-β superfamily with broad roles in embryonic patterning. In the central nervous system, BMP4 signaling is crucial for neural development: it helps establish dorsal neural tube identity, influences progenitor cell fate decisions, and regulates the balance between neuronal and glial differentiation from neural precursors [32, 79]. BMP4 also affects multiple downstream pathways (SMAD, ERK) that modulate lineage progression in neural stem cells, and its expression persists into later stages influencing glial cell behavior and responses to injury [79]. Mutations in BMP4 in humans cause pleiotropic developmental anomalies, underscoring its global morphogenetic role, but its neural activity is central to early patterning and lineage decisions in cortical and hindbrain structures. This gene-level evidence further supports one of our previous findings, suggesting a biological link between bone intensity and brain-related traits. Notably, even in this study, among the 11 newly discovered loci, variants such as rs330060 and rs6684375 show bone mineral density-related traits among the top five entries in the GWAS Catalog.

Overall, this study presents the first multimodal contrastive learning framework based on the MoCoV2 for brain MRI with genetic investigation, demonstrating that shared representations can be learned that encode common genetic and structural properties of the human brain.

## Methods

### UK Biobank

The UK Biobank [80] is a nationwide population resource encompassing approximately 500,000 participants recruited across the United Kingdom, with phenotypic, imaging, and genetic data collected between 2006 and 2010. The study operates under established ethical approval, with governance and oversight details available at https://www.ukbiobank.ac.uk/learn-more-about-uk-biobank/governance/ethics-advisory-committee. In total, 29,115 brain MRI scans were included in this study. Model development used 4,597 scans for training and 1,533 for validation. Learned representations were subsequently extracted from the remaining 22,985 individuals, corresponding to the same “discovery cohort” used in prior brain imaging GWAS analyses [5, 6]. Additional quality control excluded individuals exceeding 5 standard deviations for any UDIP, yielding final sample sizes ranging from 22,916 to 22,985 across the 128 UDIPs. Phenomics-wide association analyses (PhenoWAS) were conducted in overlapping subsets of individuals with both UDIPs and traditional IDPs, with sample sizes varying from 21,086 to 22,947 depending on the features analysed. Genome-wide association analyses (GWAS) were performed using the same discovery cohort as in prior studies [5, 6], comprising 22,799 to 22,867 individuals after filtering. Proteome-wide association analyses (ProWAS) were conducted in subsets of 1,002 to 2,898 individuals, consistent with the sample sizes reported in the prior ProWAS component of the brain-heart-eye axis study [81]. The reduced cohort size primarily reflects extensive missingness in the proteomic measurements and the limited overlap between proteomic and imaging datasets.

For genetic analyses, imputed genotype data from the UK Biobank [80] underwent standard quality control procedures. Variants were retained if they met thresholds of minor allele frequency > 0.0001 and missingness < 0.05, resulting in 8,925,988 single-nucleotide polymorphisms in the discovery cohort comprising 22,916 to 22,985 individuals.

### Model

#### Model Input Datasets and Preprocessing

UK Biobank MRIs were downloaded on October 15, 2021. UK Biobank participants had birth year between 1934 to 1971 with female to male ratio of 0.52. UK Biobank has provided a bias-field-corrected version of the brain-extracted T1-weighted (T1) and T2-weighted FLAIR (T2) images captured mainly using standard Siemens Skyra 3T running VD13A SP4 (as of October 2015), with a standard Siemens 32-channel RF receive head coil. Resolution of T1 is 1 × 1 × 1 mm, and resolution of T2 is 1.05 × 1 × 1 mm (https://biobank.ctsu.ox.ac.uk/crystal/crystal/docs/brain_mri.pdf).

To promote generalizability and minimize manual feature engineering, we adopted the standard preprocessing pipeline developed by the UK Biobank MRI team. Brain MRI preprocessing was performed primarily using FSL (https://www.fmrib.ox.ac.uk/ukbiobank/), and included defacing, brain extraction with BET, linear and non-linear registration to standard space using FLIRT and FNIRT, and bias-field correction using FAST. Bias-field-corrected, brain-extracted T1-weighted and T2-weighted FLAIR images were retained for analysis. All images were subsequently linearly registered to MNI152 space using affine transformation with 12 degrees of freedom via UK Biobank-provided precomputed matrices in FLIRT. Linear registration was chosen to normalize head size and achieve cross-subject alignment while preserving subject-specific structural deformations, in contrast to non-linear registration. The resulting linearly registered, defaced, bias-field-corrected images were used in all downstream analyses. Image intensities were normalized on a per-scan basis using Z-score normalization. Following affine registration, T1- and T2-weighted volumes had dimensions of 182 × 218 × 182 voxels and were zero-padded to 182 × 224 × 182 to enable partitioning into equal-sized patches.

To construct a high-quality deep learning dataset, we leveraged UK Biobank’s precomputed MRI quality metrics: the inverted contrast-to-noise ratio (Data Field 25735) and the discrepancy between T2 FLAIR and T1 brain images (Data Field 25736). Images with values below the 95th percentile for both metrics-corresponding to higher-quality scans-were retained. For participants with multiple visits, only the first scan was kept to ensure consistency across the dataset. This filtering resulted in a cohort of 6,130 scans from individuals of diverse ethnic backgrounds. The dataset was randomly split into a training set of 4,597 images (75%) and a validation set of 1,533 images (25%). The validation set was reserved exclusively for hyperparameter tuning and model checkpointing during training.

#### Model training

We adopt MoCo v2 (Momentum Contrast) as the contrastive learning framework [23], which consists of a query encoder and a key encoder that share the same ViT architecture and are initialized with identical weights. The key encoder is updated as an exponential moving average of the query encoder parameters, allowing it to evolve smoothly and provide stable target representations during training.

In our multimodal setting, paired T1- and T2-weighted MRI from the same subject are treated as positive views. One modality is processed by the query encoder and the corresponding paired modality is processed by the key encoder, forming cross-modality positive pairs that encourage alignment of representations across MRI modalities. This contrastive formulation is symmetric, such that both T1 and T2 can alternatively serve as the query or key view, and the final loss is computed as the average of the two directional contrastive losses.

Negative samples are obtained from a first-in-first-out queue that stores representations generated by the key encoder from previous mini-batches. All entries in the queue serve as negative samples during contrastive learning, with the oldest representations located at the front of the queue. After each training iteration, newly computed key representations are enqueued at the back of the queue, while the oldest entries are dequeued from the front to maintain a fixed queue size. This queue-based mechanism enables the use of a large and diverse set of negative samples without increasing batch size.

The model is trained using the InfoNCE contrastive loss, which encourages alignment between cross-modality positive pairs while pushing representations away from negatives in the dictionary. This multimodal contrastive design enables the model to learn modality-invariant representations of brain structure.

The model is parameterized as follows. The ViT encoder consists of 12 transformer layers with 6 attention heads per layer and an embedding dimension of 384. The projection head maps encoder outputs to a 256-dimensional contrastive embedding space. The contrastive dictionary is implemented as a queue with a size of 65,536 entries, and the momentum coefficient for updating the key encoder is set to 0.999. Cosine similarity is used in the contrastive objective with a temperature parameter of 0.07. Training is performed using the AdamW optimizer with an initial learning rate of 1 × 10⁻⁴ and a CosineAnnealingWarmRestarts learning rate scheduler (T₀ = 10, T_mult = 2, η_min = 1 × 10⁻⁶).The model was trained for 300 epochs on 4 Nvidia H100 GPUs.

#### Model embedding extraction

We froze the ViT encoder and removed the projection head, then applied the model to T1- and T2-weighted MRIs of 22,985 held-out individuals (discovery cohort). This produced representations of size 22,985 samples × 2,366 patches × 384 dimensions. To reduce dimensionality, the 2,366 × 384 patch features for each subject were compressed into 128 principal components using PCA.

#### CKA

To quantify the similarity between embeddings from different modalities or models, we performed linear Centered Kernel Alignment (CKA) [27]. First, samples containing missing values in either embedding were removed. For cross-modality comparison, paired T1- and T2-weighted MRI embeddings from the same subject formed positive pairs, while negative pairs were generated by randomly shuffling subject assignments between embeddings.

To estimate variability in the similarity measures, we used bootstrap sampling for positive pairs and repeated random permutations for negative pairs. Linear CKA was then computed on these aligned embeddings, producing a similarity score ranging from 0 (no similarity) to 1 (identical representations). The positive-pair distribution captures modality-consistent similarity, whereas the negative-pair distribution serves as a baseline expected by chance.

#### CCA

To assess the shared information captured by embeddings from paired T1- and T2-weighted MRI, we performed canonical correlation analysis (CCA). First, per-subject embeddings were loaded from CSV files for each principal component (128 features per modality) and merged by subject identifiers (IIDs). Samples containing missing values in either modality were excluded to ensure complete data alignment.

The resulting feature matrices were standardized to zero mean and unit variance prior to CCA. CCA was then applied using all possible components, with the number of canonical components set to the minimum of the number of subjects minus one and the number of features. This procedure identifies linear combinations of T1 and T2 embeddings that maximize cross-modality correlation.

Canonical correlations were computed for each component, producing a vector of correlation coefficients that quantify the strength of shared structure between the modalities. The distribution of canonical correlations was summarized using mean, maximum, and the number of components exceeding thresholds (ρ > 0.3 and ρ > 0.5) to assess the degree of alignment. Results were visualized as a bar plot of canonical correlations across components, highlighting components with high cross-modality correspondence.

#### UMAP

To visualize the structure of learned embeddings from paired T1- and T2-weighted MRI, we employed Uniform Manifold Approximation and Projection (UMAP) for dimensionality reduction. Samples with missing values in either modality were excluded, yielding a set of complete paired embeddings.

Because SM-UDIPs for T1 and T2 were generated using independently trained ViT autoencoders, their embeddings do not reside in a shared coordinate system, and direct distances between modalities are not meaningful. Therefore, for SM-UDIPs, UMAP visualization was performed on representations transformed by CCA, which explicitly maps T1 and T2 embeddings into a shared latent space. UMAP was applied to the CCA-aligned representations using two output dimensions, 30 nearest neighbors, and a minimum distance of 0.1, enabling qualitative assessment of cross-modality alignment.

In contrast, MM-UDIPs for T1 and T2 were derived from a shared encoder and thus naturally reside in a joint embedding space. For these representations, UMAP was performed both on the original MM-UDIP embeddings and on their CCA-transformed representations, using the same UMAP parameters for consistency. The resulting low-dimensional embeddings were visualized in two-dimensional scatter plots, with samples colored by modality.

### Phenome-wide associations

Phenome-wide association analyses (PhenoWAS) were performed to link UDIPs with 179 traditional IDPs (**Supp Table 6**) available in the downloaded UK Biobank dataset (22,945 < N < 22,947). Associations were tested using linear regression, with each traditional IDP as the dependent variable and the corresponding UDIP as the independent variable. Models included multiple covariates to control for potential confounding, including age (Field ID: 21003), sex (Field ID: 31), brain positioning in the scanner (lateral, transverse, and longitudinal axes; Field IDs: 25756-25758), intracranial volume (Field ID: 25000), body weight (Field ID: 21002), height (Field ID: 50), waist circumference (Field ID: 48), assessment centre (Field ID: 54), and the first 40 genetic principal components.

### Proteome-wide associations

Proteome-wide association analyses (ProWAS) were conducted using two sets of 128 UDIPs mapped to 2,923 unique proteins measured on the Olink platform (sample sizes ranging from 10,018 to 39,489). The raw data were generated and processed by the UK Biobank Pharma Proteomics Project, and we focused on the first instance of the proteomics measurements (‘instance’ = 0). To integrate metadata, Olink files containing coding information, batch identifiers, assay details, and limits of detection (LOD; Category ID: 1839) were merged with the protein dataset. Normalized Protein eXpression (NPX) values below the mean protein-specific LOD across plates were excluded, and proteins with fewer than 10,000 observations were removed. After applying these filters, 2,457 proteins remained for analysis. Matching these proteins to UDIP sample IDs yielded a final ProWAS cohort of 1,002 to 2,898 individuals, consistent with previous reports [81]. Analyses were adjusted for a range of covariates to control for potential confounding, including demographic factors (age, Field ID: 21003; sex, Field ID: 31), anthropometrics (body weight, Field ID: 21002; height, Field ID: 50; waist circumference, Field ID: 48; BMI, Field ID: 23104), brain- and imaging-specific variables (scanner positioning along lateral, transverse, and longitudinal axes, Field IDs: 25756-25758; head motion, Field ID: 25741; intracranial volume), assessment centre (Field ID: 54), and the first 40 genetic principal components.

Following the ProWAS, functional analyses were performed including Gene Ontology (GO) biological process enrichment, KEGG pathway analysis, and STRING network exploration.

Protein identifiers were converted to Entrez gene IDs using the org.Hs.eg.db (v3.17.0) annotation database in R, excluding proteins without valid IDs. GO biological process enrichment was performed using clusterProfiler (v4.8.0) with multiple testing correction by the Benjamini-Hochberg method (p.adjust < 0.05). Significant GO terms were visualized with bar plots displaying the number of associated proteins.

KEGG pathway enrichment was conducted using clusterProfiler (enrichKEGG, organism = “hsa”, p < 0.05), with pathway names validated through the KEGG REST API via httr (v1.4.7) for datasets without clusterProfiler. Results were saved as CSV files and visualized with bar plots highlighting the top pathways. All analyses were performed in R (≥4.3.0) using CRAN packages ggplot2 (v3.4.3), dplyr (v1.1.3), pheatmap (v1.0.12), and igraph (v1.4.6). Visualizations facilitated the interpretation of enriched biological processes and pathways for the protein datasets.

Protein-protein interaction networks were constructed using the STRING database (v11.5) by querying the list of proteins of interest with a high-confidence interaction score threshold (≥700) for Homo sapiens. The resulting interaction data, including protein pairs and combined confidence scores, were imported into Python and visualized as undirected weighted networks using igraph (v1.4.6). Nodes represent proteins, with size proportional to the number of interactions (degree), and edges represent interactions, weighted by STRING confidence scores. Proteins were optionally colored based on functional categories derived from Gene Ontology biological process annotations. Final network figures highlighted hub proteins and overall network topology for downstream functional interpretation.

### Genetic analyses

We implemented a comprehensive quality-control pipeline on the downloaded genetic data, following our previous work [14], resulting in 8,469,833 single-nucleotide polymorphisms (SNPs) across 22,985 individuals of European ancestry. Briefly, duplicated variants across the 22 autosomal chromosomes were removed, and individuals whose genetically inferred sex did not match their self-reported sex were excluded. Additional filtering criteria removed variants with a minor allele frequency (MAF) below 0.01% or a genotyping missing rate exceeding 5%.

#### GWAS

GWAS were conducted on 22,867 individuals for 128 extracted UDIPs. Associations between SNPs and UDIPs were tested using linear mixed models implemented in FastGWA [25] within GCTA (v1.94.1), with a minor allele frequency threshold of 0.01. Covariates included age (Field ID: 21003), age², sex (Field ID: 31), sex × age, sex × age², the first 10 genetic principal components (Field ID: 22009), intracranial volume (Field ID: 25000), inverted contrast-to-noise ratio (Field ID: 25735), head positioning in the scanner (Field IDs: 25756-25758), scanner table position (Field ID: 25759), assessment center location (Field ID: 54), and date of assessment (Field ID: 53). For participants with multiple visits, only the first scan was retained. A Bonferroni-corrected significance threshold of 5 × 10⁻⁸/128 was applied, and summary statistics were compiled with the most significant p-value (minP) reported for each SNP.

#### Annotation of genomic loci

Genomic loci were annotated using FUMA [26]. Lead SNPs were first identified based on linkage disequilibrium (r² ≤ 0.1) and physical proximity (<250 kb), and assigned to non-overlapping genomic loci. Within each locus, the SNP with the lowest p-value (the top lead SNP) was used to represent the locus.

#### Overlap of genomic loci

To assess locus overlap, each identified locus was extended by 125 kb on either side, and interval trees were constructed for each chromosome. Loci from our analysis were then queried against these interval trees to determine overlapping regions.

#### SNP-based heritability

SNP-based heritability (h²) was estimated using GCTA, which constructs a genetic relationship matrix from raw genotype data, providing a robust estimate of the proportion of phenotypic variance explained by common SNPs and helping to address the problem of ‘missing heritability’ [37].

#### Genetic correlation

Genetic correlations (gc) between T1- and T2-derived UDIPs were estimated using LDSC. Precomputed linkage disequilibrium scores from the 1000 Genomes European reference panel were used, while all other parameters were kept at their default settings. Significance thresholds were adjusted using Bonferroni correction, based on the number of UDIPs analyzed (N = 128).

#### Gene-drug-disease network

To map potential gene-drug-disease relationships, we first identified genes significantly associated with the 128 T1- and T2-derived brain UDIPs. SNPs were mapped to genes based on their physical positions using the Phase 3 1000 Genomes reference panel for European ancestry, and gene-level association tests were performed with MAGMA [82] using SNP GWAS summary statistics. Bonferroni correction (p < 0.05/18,943) was applied to define significant genes, yielding 1,376 genes for T1 and 1,341 genes for T2. These genes were then analyzed with GREP [38] to test for enrichment in drug-target sets curated from DrugBank [39], corresponding to specific clinical indication categories (ICD codes). Fisher’s exact tests assessed overrepresentation, and multiple testing was corrected using the false discovery rate, allowing identification of statistically significant gene-drug-disease associations.

### Disease and cognition prediction

We assessed the ability of the 128 combined T1- and T2-derived UDIPs to predict 2 systemic disease categories as well as 8 cognitive performance measures. 2 systemic disease categories include Mental and behavioural disorders (F) and Nervous system diseases (G). 8 cognitive scores include Matrix Pattern Completion: Assessed via the number of puzzles solved correctly (Field ID: 21004); Digit Symbol Substitution Test (DSST): Assessed via the number of correct symbol-digit matches (Field ID: 23324); Tower Rearranging: Measured by the number of puzzles correctly solved (Field ID: 6373); Trail Making Test A (TMT-1): Measured by the duration to complete a numeric path (Field ID: 6348); Trail Making Test B (TMT-2): Measured by the duration to complete an alphanumeric path (Field ID: 6350); Fluid Intelligence: Derived from the fluid intelligence score (Field ID: 20016); Numeric Memory: Assessed via the maximum number of digits remembered correctly (Field ID: 4282); Reaction Time: Measured as the mean time to correctly identify matches (Field ID: 20023).

Patient cohorts for the two disease categories were defined using ICD-10 codes available through UK Biobank (Data Fields: 41270 and 41202), while the healthy control group comprised individuals with no ICD-10-based diagnoses. The overlapping system-disease dataset included 1,851 participants with mental and behavioural disorders (Category F) and 2,333 participants with nervous system disorders (Category G). Cognitive score data were available for a larger subset, encompassing 15,571 to 21,349 participants, depending on the specific cognitive assessment.

#### Systemic diseases classification

We assessed the predictive power of two distinct feature sets-128 T1- and T2-derived UDIPs obtained from either single-modality (SM) or multimodal (MM) sources-using logistic regression framework combined with a nested cross-validation (CV) strategy. The evaluation leveraged separate training, validation, and test datasets. Predictive accuracy was quantified using the balanced accuracy from the nested CV, addressing potential class imbalance. The nested CV procedure consisted of an outer 3-fold CV loop and an inner 5-fold CV loop. Within each inner iteration, 67% of the training/validation data were further partitioned into 5 folds to optimize hyperparameters, enabling rigorous tuning while preventing information leakage from the test sets. We majorly report the logistic regression results as it shows better performance (**Supp Fig 8**).

#### Cognitive scores regression

Both MM-UDIPs and SM-UDIPs were used to predict eight cognitive scores using support vector regression and LASSO regression. Model performance was primarily evaluated using Pearson’s correlation coefficient (r) between predicted and observed scores in the training and test datasets. We focus on support vector regression results in the main text, as it consistently outperformed LASSO (**Supp Fig 9**).

## Data Availability

The model codes and pre-trained model checkpoints are publicly accessible at Github at https://github.com/ZhiGroup/UDIP-MoCoV2. The UDIPs and GWAS summary statistics are available upon reasonable request after removing sensitive UKbiobank patient information. Our study used data generated by the STRING data (https://string-db.org/). Genomic loci annotation used data from FUMA (https://fuma.ctglab.nl/). PheWAS used traditional IDP list from the Big40 IDP (https://open.oxcin.ox.ac.uk/ukbiobank/big40/). Individual data from UKBB can be requested with proper registration at https://www.ukbiobank.ac.uk/. The gene-drug-disease network used data from the DrugBank database (v.5.1.9; https://go.drugbank.com/). All unrestricted data supporting the findings are also available from the corresponding author upon request.

## Code Availability

The software and resources used in this study are all publicly available:

- Model: https://github.com/ZhiGroup/UDIP-MoCoV2.
- FUMA (v.1.5.0): https://fuma.ctglab.nl/, gene mapping, genomic locus annotation
- GCTA (v.1.94.1): https://yanglab.westlake.edu.cn/software/gcta/#Overview, heritability estimates and fastGWA

## Acknowledgements

This work was supported by grants from the National Institute on Aging U01AG070112 and R01AG081398.

We used a large language model to assist with language editing and clarity of presentation. The authors reviewed and edited all content and take full responsibility for the accuracy and integrity of the work.

## Ethics declarations

### Competing interests

The authors declare no conflict of interest.

### Ethical Approval

Our analysis was approved by the UTHealth Houston committee for the protection of human subjects under No. HSC-SBMI-20-1323. UKBB has secured informed consent from the participants in the use of their data for approved research projects. UKBB data was accessed via approved project 24247.

## Supplementary Information

We have included the following supplementary materials.

- Supplementary Notes
- Supplementary Figures 1-9
- Supplementary Tables 1-6

## Notes

### Competing Interest Statement

The authors have declared no competing interest.

